# Localization of a feline autosomal dominant dwarfism locus: a novel model of chondrodysplasia

**DOI:** 10.1101/687210

**Authors:** L.A. Lyons, D.B. Fox, K.L. Chesney, L.G. Britt, R.M. Buckley, J.R. Coates, B. Gandolfi, R.A. Grahn, M.J. Hamilton, J.R. Middleton, S.T. Sellers, N.A. Villani, S. Pfleuger, the 99 Lives Consortium

## Abstract

Despite the contribution of a few major genes for disproportionate dwarfism in humans, many dwarf patients are yet genetically undiagnosed. In domestic cats, disproportionate dwarfism has led to the development of a defined breed, the Munchkin or Minuet. This study examined the genetic aspects of feline dwarfism to consider cats as a new biomedical model. DNA from dwarf cats was genetically analyzed using parentage, linkage, and genome-wide association studies as well as whole genome sequencing. Each genetic approach localized the dwarfism phenotype to a region on cat chromosome B1. No coding variants suspected as causal for the feline dwarfism were identified but a critical region of ∼5.7 Mb from B1:170,278,183-175,975,857 was defined, which implicates a novel gene controlling disproportionate dwarfism. A yet unidentified but novel gene variant, likely structural or regulatory, produces disproportionate dwarfism in cats, which may define undiagnosed human patients.

## Introduction

Dwarfism is a genetic and or endocrine abnormality causing an animal to be less than normal size and lacking the capacity for normal growth. In humans, any person less than 147 cm for adult height is considered a dwarf^1^. Two general categories of dwarfism are defined, disproportionate and proportionate dwarfism. Disproportionate dwarfs possess shortened limbs with a normal trunk while proportionate dwarfs are uniformly small^2,3^. Disproportionate dwarfs result from genetic disorders involving bone and or cartilage, while the majority of instances of proportionate dwarfism result from hormonal abnormalities^4,5^.

In humans, over 200 different forms of dwarfism are defined and approximately 1 in 20,000 newborn infants display congenitally shortened limbs, i.e. disproportionate dwarfism. DNA variants in one gene, *fibroblast growth factor receptor 3 (FGFR3)*, account for a majority of human disproportionate dwarfism with hypochondrodysplasia (HCH) (OMIM <u>#146000</u>) or achondroplasia (OMIM <u>#100800</u>) constituting the majority of diagnosed cases. The *FGFR3* G1138M variant accounts for 99% of all ACH^6–8^ while an estimated 70% of HCHs result from variants in *FGFR3* (see reviews^9,10^). However, many forms of disproportionate human dwarfism are still undefined.

Feline disproportionate dwarfism (OMIA 000299-9685) has only briefly been mentioned in the scientific literature and has not been clinically characterized^11^. Breeds of cat have developed from disproportionate dwarf cats, termed Munchkins, Napoleons or Minuets (**Fig. 1**)^12^. The cats are at least mesomelic, with shortened forelimbs and hind limbs, implying the humerus, the radius and ulna and the femur, the tibia and fibula have deficient growth. Pre-mature ossification of epiphyseal plate cartilage is unknown. As reported by breeders, the cats have no indication of gender bias in the deformity and no indication of homozygotes. The feline phenotype lacks the frontal bossing and other maladies associated with achondroplasia^13^, although the clinical features of the dwarf cats have not been extensively defined. An autosomal dominant mode of inheritance has been suggested. Thus, feline dwarfism closely parallels the human HCH phenotype, although the genetic cause of feline dwarfism is undetermined. The cat may be an important biomedical model for human chondrodystrophic disorders and potentially osteoarthrosis and degenerative joint disease.

**Figure 1.**
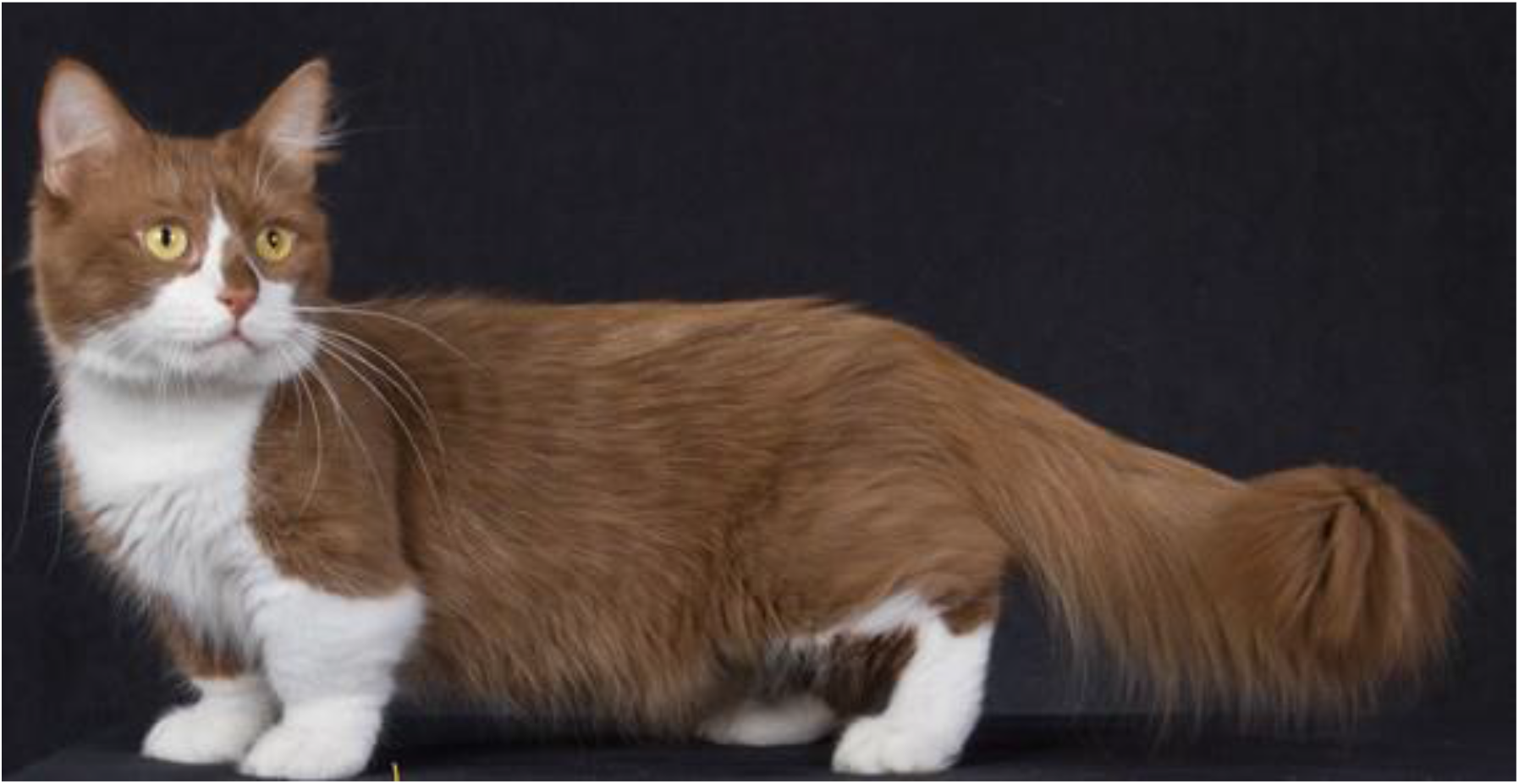
Dwarfism in domestic cats. Identified in random bred cats, dwarfism in cats presents as varying degrees of shorted long bones. The random bred cats have been bred to Persians and other breeds and have various breed names, such as, Napoleons, Minuets or Munchkins. Although the breed is controversial due to stature, additional adverse health problems have not been documented. A variety of hair coats and pelage colorations are accepted in the breed standard in The International Cat Association (TICA). Image courtesy of Terri Harris.

To genetically define the disproportionate dwarfism in the domestic cats and implicate candidate genes for this biomedical model, genetic studies were conducted in dwarf cats. Linkage analyses and genome-wide association localized the dwarfism phenotype to a region on cat chromosome B1, which has homology to human chromosome 4 but exclusive of the region of *FGFR3*, supporting the plausibility of cat disproportionate dwarfism as a novel biomedical model to help identify the approximately 30% of HCH human cases that do not yet have gene associated mutations.

## Materials and Methods

### DNA analyses - pedigree and DNA extraction

Buccal swabs or EDTA anti-coagulated whole blood samples from cats were solicited from owners of dwarf cats for DNA isolation over a period of twenty years. The familial relationships and dwarfism phenotypes were reported by the owners and breeders. DNA was isolated using the Qiagen DNAeasy kit according to the manufacturers protocol (Qiagen, Inc. Hilden, Germany) or standard organic extraction. All sample collections were conducted in accordance with an approved University of California, Davis Institutional Animal Care and Use protocols 11977, 15117, and 16691 and University of Missouri protocols 7808 and 8292.

### DNA Analyses – Short Tandem Repeat (STR) Analysis

To develop a pedigree, parentage of the cats were verified with a panel of feline-derived short tandem repeats (STRs) as previously described^14^. STR fragment sizes were determined using STRand analysis software^15^.

### DNA Analyses – Linkage Analysis

Two-point linkage between the STRs and the dwarfism phenotype was conducted as previously described^16^ using the LINKAGE and FASTLINK software programs^17–19^. Loops in the pedigree were broken by duplication of individuals. The dwarfism phenotype was coded as a fully penetrant autosomal dominant trait with non-variable expression. All matings were between a dwarf and a non-dwarf cat. No homozygous cats were predicted by the breedings nor have been reported by breeders, thus dwarf cats were assumed to be heterozygous. STR positions were determined by aligning the unique sequence flanking of the repeats to cat reference genome Felis_Catus_9.0 (GCF_000181335.3/).

### DNA Analyses – Genome-wide association

The initial dataset for the SNP array genome-wide analysis comprised 95 cats, including 26 affected dwarf cases that were unrelated to the third generation from provided pedigree information, 11 related non-dwarf cats, and cats from the closely related breeds, Scottish fold (n = 34), British shorthair (n = 14) and Selkirk rex (n = 10) for controls^20^. Approximately 600 ng of genomic DNA was submitted to Neogene, Inc (Lincoln, NE, USA) for genotyping on the Illumina Infinium Feline 63K iSelect DNA array (Illumina, Inc., San Diego, CA). Genotyping and analysis was performed as previously described^21^. SNP genotyping rate and minor allele frequency was evaluated using PLINK^22^. SNPs with a MAF < 5%, genotyping rate < 90%, and individuals genotyped for < 90% of SNPs were excluded from downstream analyses. Inflation of p-values was Λ evaluated by calculating the genomic inflation factor (λ). The 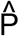 for each individual, an identity by descent analysis, and multidimensional scaling (MDS) was calculated using PLINK to evaluate population substructure within cases and controls (data not shown).

To reduce λ, cats not tightly clustered and/or highly related with a 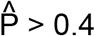 were removed from downstream analyses. Selection for each case to the closest control using the values from the MDS dimensions was attempted.

### Whole genome sequencing

Three unrelated dwarf cats were submitted for whole genome sequencing as part of the 99 Lives Cat Genome Sequencing Consortium as previously described^23–26^ (BioProject PRJNA308208, PRJNA288177; BioSamples: SAMN05980349, SAMN05980348, SAMN05980352). Reads were mapped to Felis_catus_9.0 (GCF_000181335.3/) and assigned to read groups using BWA-MEM from Burrows-Wheeler Aligner version 0.7.17^27^. Duplicate reads were marked using MarkDuplicates from Picard tools version 2.1.1 (http://broadinstitute.github.io/picard/), with OPTICAL_DUPLICATE_PIXEL_DISTANCE set at 2500. Genome Analysis Toolkit version 3.8 (GATK 3.8) were used to further process the sequence data^28^. Indel realignment was performed with RealignerTargetCreator and IndelRealigner^29^ and SNPs, and Indels were called using HaplotypeCaller in gVCF mode (-ERC GVCF)^30^. The gVCFs were combined into groups of ∼20 individuals using CombineGVCFs and were genotyped simultaneously using GenotypeGVCFs. Throughout, Samtools version 1.7 sort, index, view, and cat functions were used to process BAM files between individual tasks^31^. Together these processes produced a single VCF comprised of 195 cats for downstream analysis. DNA variants were viewed, filtered and annotated using VarSeq (Golden Helix, Boseman, MT) with the Ensembl Release 94 Felis_catus_9.0 genome annotation^32^. For variant prioritization, candidate variants were required to be within the region implicated by the GWAS (chr B1 170 – 187 Mb), heterozygous in all three dwarf cats, and absent in 71 normal cats also in the 99 Lives dataset (PRJNA308208).

## Results

### Pedigree Analysis

The dwarfism phenotype in the affected Munchkin cats is unique and distinct, with only mild variations in presentation, therefore diagnosis of affected kittens is not confounded by other congenital birth defects in cats (**Fig 1**). A six-generation, 83-member pedigree (65 informative meioses) was created based upon owner supplied pedigrees, parentage analyses and phenotypes (**Fig. 2**). The parentage analysis confirmed reported relationships of the cats (data not shown). Sires and dams bred into the pedigree were, in general, of random bred stock and were not related. All matings and all dwarf offspring in the pedigree had at least one dwarf parent. Complete litter data was available for nine matings, one was a breeding of two dwarf cats that produced six kittens, including one dwarf male kitten, three normal male kittens and two normal female kittens. Eight matings had one dwarf parent, producing eight of 19 dwarf male kittens and seven of 17 dwarf female kittens. Matings that had complete litter information showed no sex bias with 39% of males and 37% of females displaying the dwarf phenotype. No litter produced all dwarf kittens, thus the dwarf parents are expected to be heterozygous. A heterozygous affected by normal mating would predict 50% affected offspring, 42% of progeny displayed the dwarf phenotype in the pedigree. For the 10 matings with both parents having dwarfism, none of the Munchkin offspring (n = 6) are homozygous for the STR in complete linkage with the dwarf phenotype. Also, normal sized cats were produced from these matings, further excluding a recessive mode of inheritance.

**Figure 2.**
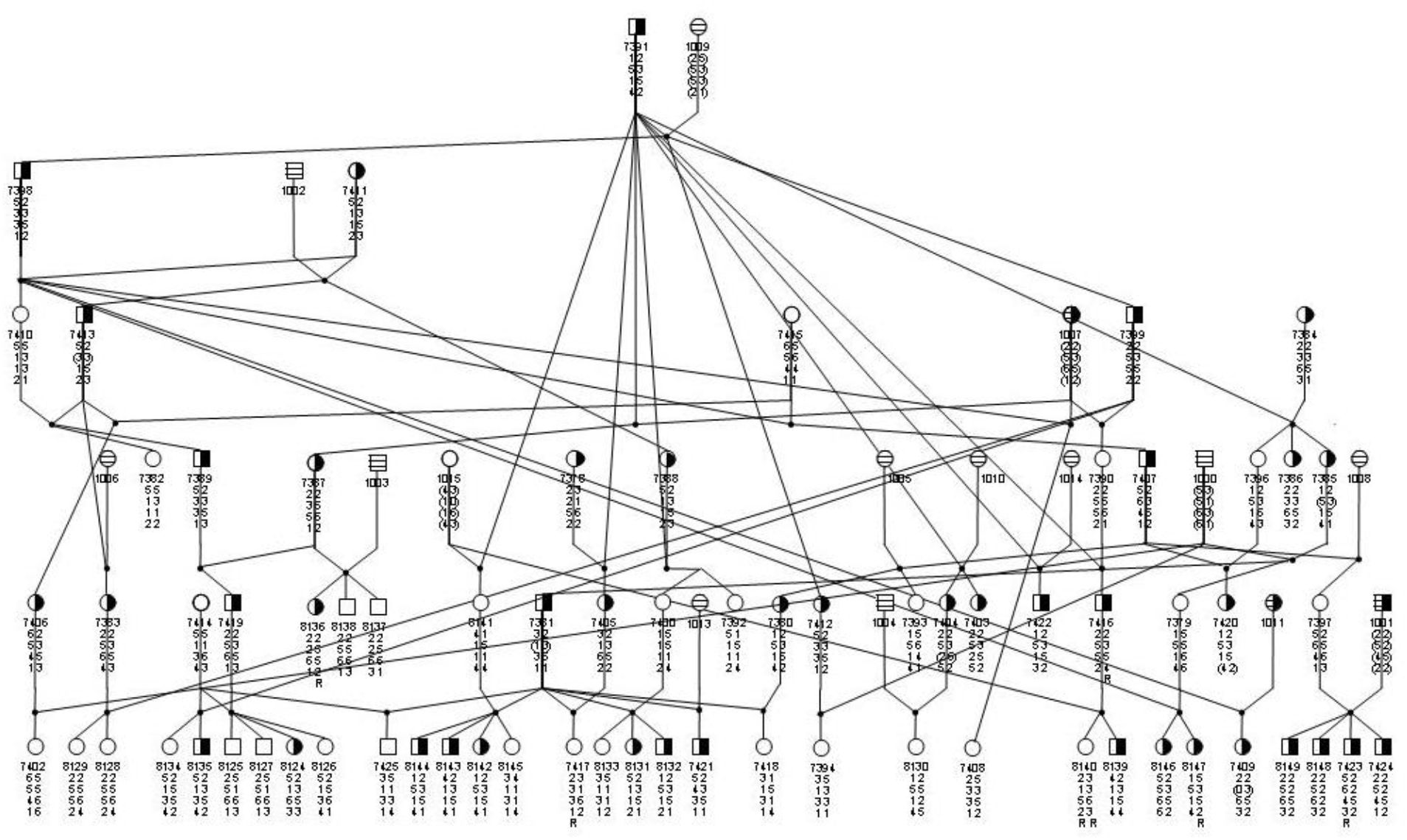
Six Generation 83-member pedigree segregating for the dwarf phenotype. Squares represent males, circles females. Open shapes are wildtype, half-filled symbols are phenotypically dwarf cats. Cross hatched symbols were not available for typing. Four-digit numbers under each symbol are animal ID numbers. Alleles for STRs *FCA149, FCA827* and *FCA152* are under each identifier. Inferred types are in parenthesis. Recombinants (N = 6) are indicated by R.

### Linkage analysis

One STR genotyped to confirm parentage, *FCA149*, indicated significant linkage (Z = 5.43. θ = 0.05) to the dwarfism phenotype and is located on feline chromosome B1^33^. Additional publicly available regional STRs (n = 8)^34–36^ were genotyped to refine the linkage region (**Table 1**). Marker *FCA827* was completely linked to the dwarfism phenotype, Z = 15.05, θ = 0.00, which is positioned at B1:174,566,680 – 174,566,701. Recombinants were detected with flanking markers *FCA149* and *FCA152*, defining a minimal critical region from B1:167,386,513 – 179,961,345, a 12 Mb as based on STR positions on the cat genome assembly.

**Table 1:** Chromosome B1 STRs with linkage to feline dwarfism.

| **STR** | **LOD (Z)** | **Theta (Θ)** | **Start** | **End** |
| --- | --- | --- | --- | --- |
| FCA820 | 0.03 | 0.40 | chrB1:97271192 | chrB1:97271050 |
| FCA730 | 1.42 | 0.20 | chrB1:182979793 | chrB1:182979657 |
| FCA097 | 0.39 | 0.30 | chrB1:154184908 | chrB1:154184757 |
| FCA257 | 0.22 | 0.40 | chrB1:158429698 | chrB1:158429838 |
| FCA149 | 5.43 | 0.05 | chrB1:167386536 | chrB1:167386414 |
| **FCA827** | **10.34** | **0.00** | chrB1:174566701 | chrB1:174566475 |
| FCA152 | 6.22 | 0.05 | chrB1:179961324 | chrB1:179961465 |
| FCA826 | 1.84 | 0.10 | chrB1:168743712 | chrB1:168743712 |
| FCA830 | unlinked | 0.40 | chrB1:200676672 | chrB1:200676788 |

\*Positions determined from cat genome assembly Felis\_Catus\_9.0 (GCF\_000181335.3/).

### Genome-wide association

A genome-wide case-control study was conducted on 26 cases and 50 controls. All samples passed genotype rate (MIND > 0.2) and 49,424 SNPs were analyzed after missingness (GENO > 0.2) and the frequency test (MAF < 0.05). The genomic inflation (λ) was 2.95, suggesting inflation is due to the population substructure as suggested by the MDS (**Fig. 4**). The highest significant association was identified on cat chromosome B1 at position 179,190,128 (**Fig. 3**). The P-values and positions of the 36 most associated SNPs are presented in **Table 2**. A majority of the highly associated SNPs reside on cat chromosome B1, from position 170,786,914 - 186,605,881, a ∼16 Mb critical region. Considering the linkage boundary of STR *FCA152* (B1:179,961,345), the critical region reduces to ∼ 9.2 Mb B1:170,786,914 – 179,961,345.

**Figure 3.**
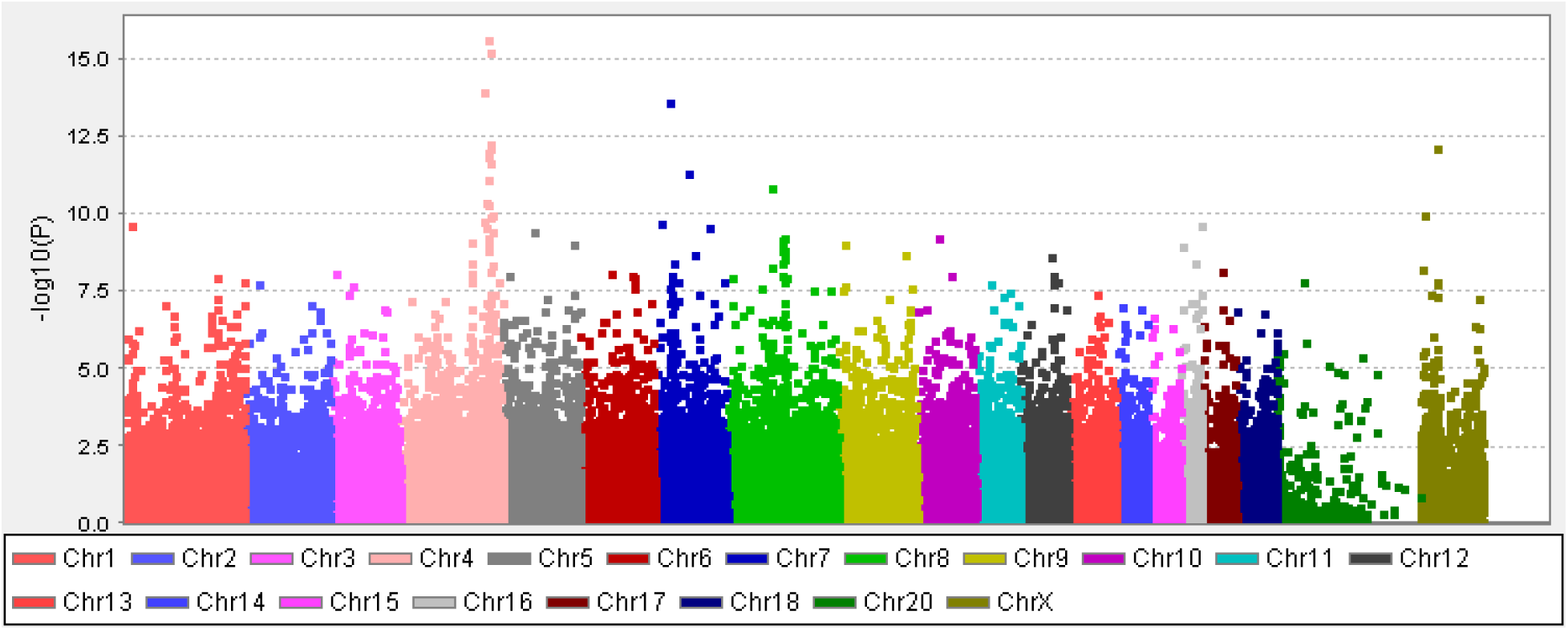
Manhattan plot of genome-wide association for feline dwarfism. Manhattan plots of the Munchkin GWAS. The plot represents the *P_raw_* values of each SNP included in the case-control association study. The association study compared the Munchkin cats to non-standard Munchkin cats, British shorthair, Selkirk rex and Scottish fold. A significant association was found with chromosome B1. Over 20 SNPs show association on cat chromosome B1 from 170 – 186 Mb. Chromosome 4 corresponds to cat chromosome B1.

**Figure 4.**
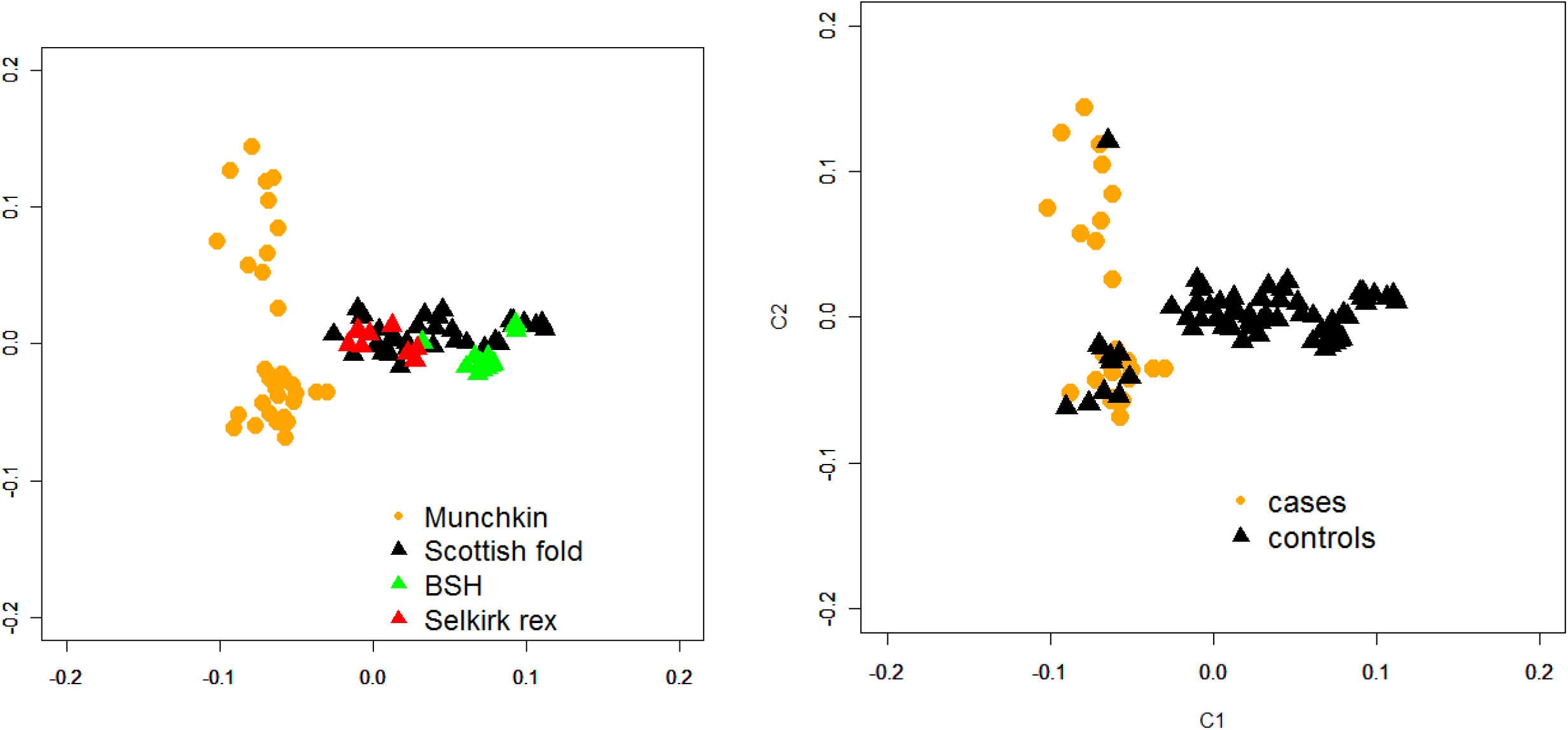
Multi-dimensional scaling (MDS) in two dimensions of samples that passed quality control for dwarf cat analysis. Left) All cats, all breeds in the preliminary analysis. Right) MDS of cases and controls included in the analysis, indicating substructure of the cats.

**Table 2:**
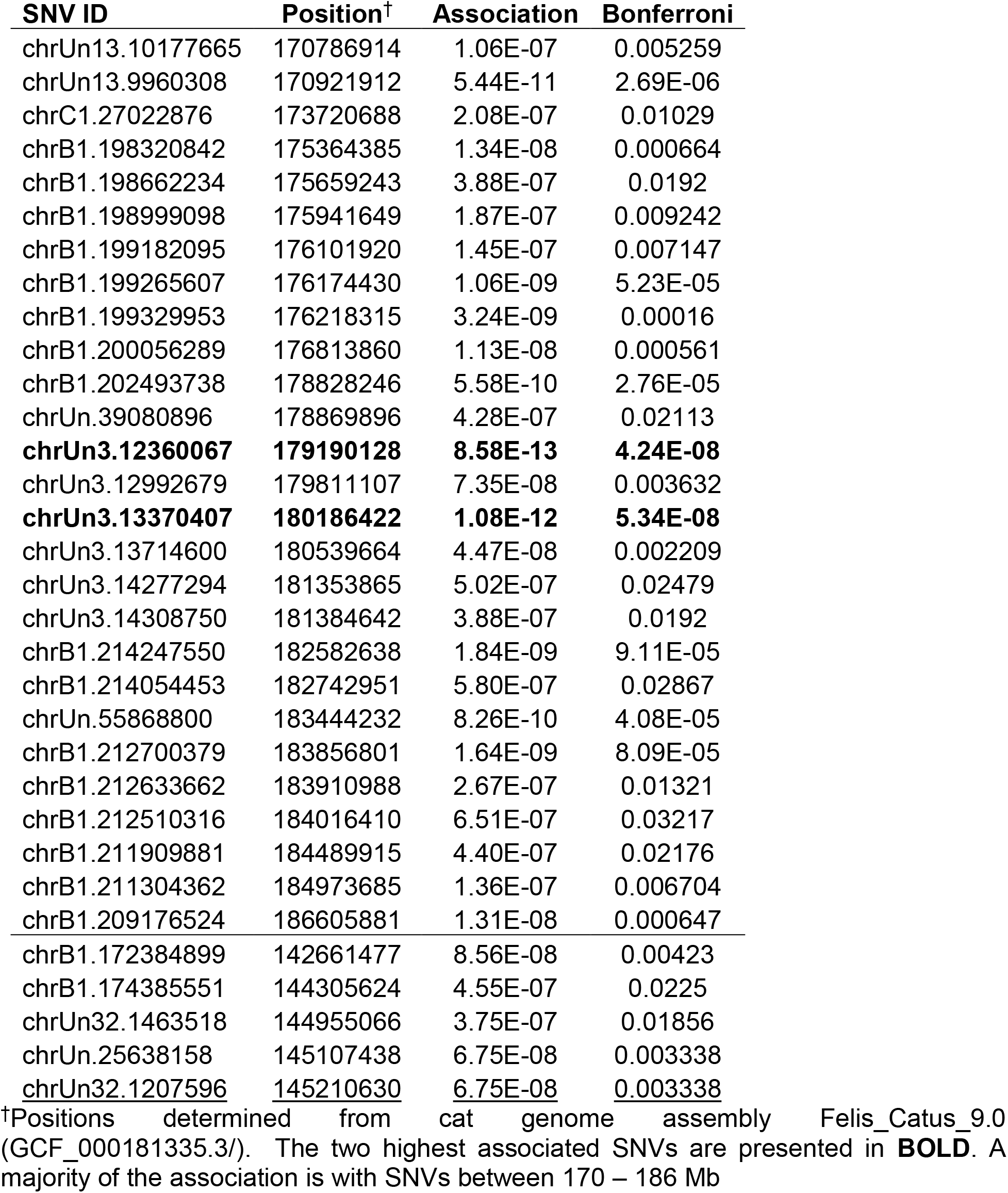
Chromosome B1 SNVs associated with feline dwarfism.

| **SNV ID** | **Position<sup>†</sup>** | **Association** | **Bonferroni** |
| --- | --- | --- | --- |
| chrUn13.10177665 | 170786914 | 1.06E-07 | 0.005259 |
| chrUn13.9960308 | 170921912 | 5.44E-11 | 2.69E-06 |
| chrC1.27022876 | 173720688 | 2.08E-07 | 0.01029 |
| chrB1.198320842 | 175364385 | 1.34E-08 | 0.000664 |
| chrB1.198662234 | 175659243 | 3.88E-07 | 0.0192 |
| chrB1.198999098 | 175941649 | 1.87E-07 | 0.009242 |
| chrB1.199182095 | 176101920 | 1.45E-07 | 0.007147 |
| chrB1.199265607 | 176174430 | 1.06E-09 | 5.23E-05 |
| chrB1.199329953 | 176218315 | 3.24E-09 | 0.00016 |
| chrB1.200056289 | 176813860 | 1.13E-08 | 0.000561 |
| chrB1.202493738 | 178828246 | 5.58E-10 | 2.76E-05 |
| chrUn.39080896 | 178869896 | 4.28E-07 | 0.02113 |
| **chrUn3.12360067** | **179190128** | **8.58E-13** | **4.24E-08** |
| chrUn3.12992679 | 179811107 | 7.35E-08 | 0.003632 |
| **chrUn3.13370407** | **180186422** | **1.08E-12** | **5.34E-08** |
| chrUn3.13714600 | 180539664 | 4.47E-08 | 0.002209 |
| chrUn3.14277294 | 181353865 | 5.02E-07 | 0.02479 |
| chrUn3.14308750 | 181384642 | 3.88E-07 | 0.0192 |
| chrB1.214247550 | 182582638 | 1.84E-09 | 9.11E-05 |
| chrB1.214054453 | 182742951 | 5.80E-07 | 0.02867 |
| chrUn.55868800 | 183444232 | 8.26E-10 | 4.08E-05 |
| chrB1.212700379 | 183856801 | 1.64E-09 | 8.09E-05 |
| chrB1.212633662 | 183910988 | 2.67E-07 | 0.01321 |
| chrB1.212510316 | 184016410 | 6.51E-07 | 0.03217 |
| chrB1.211909881 | 184489915 | 4.40E-07 | 0.02176 |
| chrB1.211304362 | 184973685 | 1.36E-07 | 0.006704 |
| chrB1.209176524 | 186605881 | 1.31E-08 | 0.000647 |
| chrB1.172384899 | 142661477 | 8.56E-08 | 0.00423 |
| chrB1.174385551 | 144305624 | 4.55E-07 | 0.0225 |
| chrUn32.1463518 | 144955066 | 3.75E-07 | 0.01856 |
| chrUn.25638158 | 145107438 | 6.75E-08 | 0.003338 |
| <u>chrUn32.1207596</u> | <u>145210630</u> | <u>6.75E-08</u> | <u>0.003338</u> |

<sup>†</sup>Positions determined from cat genome assembly Felis\_Catus\_9.0 (GCF\_000181335.3/). The two highest associated SNVs are presented in **BOLD**. A majority of the association is with SNVs between 170 – 186 Mb

### Whole genome sequencing

Approximately 30x genome coverage was produced from the three dwarf cats. An ∼17 Mb region containing 369,708 variants was examined. Only 65 candidate variants were identified after considering the dwarf cats to be heterozygous and the variant absent in the 192 additional cats in the 99 Lives dataset. However, no variants were coding nor protein altering, including 33 intergenic and 32 intronic variants found within the eight genes, including, *LIMCH1, APBB2, RBM47, CHRNA9, UGDH, RFC1, WDR19*, and *TMEM156* (**STable 1**). These variants encompassed the 5.7 Mb region of B1:170,278,183 - 175,975,857, further refining the critical region to B1:170,786,914 – 175,975,857, ∼5.2 Mb. Overall, 48 genes and transcripts are defined within this critical region (**STable 2**).

## Discussion

The domestic cat is one of the few species with an autosomal dominant mode of inheritance for dwarfism that does not have other syndromic features, a strong model for HCH. The Munchkin cat is a relatively new breed of domestic cat and has developed from random bred dwarf cats. A random bred, short-limbed pregnant female was found in Louisiana in 1983^37,38^. The resulting litter also possessed short-limbed cats, one of which was used as a breeding male from which all Munchkins are supposedly derived. This recent de novo mutation could explain some of the high genomic inflation in the GWAS. Other spontaneous dwarf cats have been reported in the lay literature, but their contribution in the Munchkin breeding program is speculative. Within ten years, the breed has sufficiently grown in popularity to be accepted for championship show status by The International Cat Association (TICA) in 1994^31^, but is banned from registration in many other cat fancy registries, thus, is mainly bred in the USA. The segregation in the pedigree suggested an autosomal dominant mode of inheritance for feline dwarfism in the Munchkin cat breed. No individuals from six matings of two dwarf parents were homozygous for the linked marker, which suggests lethality *in utero*. An autosomal recessive mode of inheritance was also eliminated as two dwarf parents produced offspring with normal limb length.

Nearly 4000 human infants are born each year with some form of dwarfism. Disproportionate dwarfism constitutes the vast majority of congenital dwarfism and has both sex-linked and autosomal dominant modes of inheritance. The cat autosomal dominant disproportionate dwarfism most closely resembles human HCH as the cat has no noted health concerns other than the limb abnormalities. However, the genetic studies did not implicate *FGFR3* in cats and Sanger sequencing of the coding region of *FGFR3* (data not shown) did not identify any associated variants. In dogs, *FGF4* retrogene is responsible for autosomal recessive chondroplasia in 19 canine dwarf breeds^39^. Additional variants in canine homologs for *Cartilage specific integrin alpha 10 (ITGA10)^40^, collagen alpha-2(XI) chain (COL11A1)*^41^, and two *collagen type IX* genes *(COL9A2* and *COL9A3*)^42^ cause an autosomal recessive chondrodysplasia affecting the Norwegian Elkhound and Karelian Bear Dog breeds, a mild form of disproportionate dwarfism in Labrador Retrievers, which is not associated with any obvious health problems such as secondary arthrosis, and an oculoskeletal dysplasia in Labrador retrievers and Samoyed dogs, respectively. These canine dwarfism genes are not on a chromosome that shares synteny with cat chromosome B1.

Three complementary and independent genetic analyses, including a familial-based linkage analysis, a GWAS and whole genome sequencing refined a critical region to ∼5.7 Mb from B1:170,278,183-175,975,857, which has no genes causal for dwarfism and thereby implicating a novel gene for dwarfism. Although on the same chromosome, *FGFR3* (B1:207161233 - 207179684) is not near this area of association. At least 46 genes were annotated within the critical region for feline dwarfism. Unfortunately, no coding or splice variants were identified within these genes in the region. One gene, *replication factor C, subunit 1* (*RFC1*) has a pathway associated with Hutchinson-Gilford Progeria Syndrome (OMIM: 176670), which is a rare disorder characterized by short stature, low body weight, early loss of hair, lipodystrophy, scleroderma, decreased joint mobility, osteolysis, and facial features that resemble aged persons^43^. As for all WGS-based approaches, the analyses were limited by annotation, correct genome assembly, and the ability to identify more complex variants, such as structural variants and variants that influence gene regulation. Therefore, structural or regulatory variants should be examined within these regions to further implicate one of the regional candidate genes or unannotated transcripts.

Surveillance for animal welfare concerns resulting from pedigree animal breeding practices has increased since the BBC documentary *Pedigree Dogs Exposed (2008)*^44^. Although cats do not have the same extreme conformational ranges as dogs, heightened scrutiny of breeding practices has extended to cats. Orthopedic diseases are commonly associated with dog breed conformation, but far less so with domestic cat breeds^45,46^. Hip dysplasia is a concern of the Maine Coon breed^47^, osteochrondrodysplasia is an inherent condition of the Scottish Fold breed, and osteoarthritis, particularly of the elbow, is now recognized as a common condition of aged cats^48,49^. The Scottish Fold, which has varying degrees of osteochondrodysplasia, has been banned from several cat registries. The Manx, which can suffer from various conditions resulting from sacral and caudal vertebral agenesis, has heightened the concern for acceptance of other cat breeds with tail abnormalities. Thus, welfare concerns have encircled the acceptance of chondrodysplastic cats as a recognized breed.

Controversy regarding the immediate and long-term health of the chondrodysplastic cats focuses on the potential for impaired ambulation, secondary osteoarthritis^50^ and intervertebral disc disease^51,52^, which is common to many chondrodysplastic dog breeds. Biapical deformities are common in dogs with limb deformities, particularly chondrodystrophic dogs^53^. *Fibroblast growth factor 4 (fgf4*) is strongly associated with autosomal recessive chondrodysplasia in at least 19 dog breeds including dachshund, corgi, and basset hound^39^. The *FGF4* variant has also been shown to be the leading cause for intervertebral disc degeneration in the chondrodysplastic dogs^54^, thus, since the cats do not have this variant, disc disease is a less likely concern. However, poor breeding practices, such as striving for the shortest legs or longest body, could lead to similar health concerns in the cats to those that plague the dwarf dog breeds^55^.

Although many chondrodysplastic dog breeds have the *FGF4* variant, selection by breeders can lead to longer straighter legs as found in several chondrodysplastic dog breeds or the severely dismorphic limbs of basset hounds. The shortest documented cat stands at 5.25 in (13.34 cm), while other cat breeds, measured from ground to withers, range from 8 – 14 in (20.32 – 35.56) (LA Lyons, personal communication). Selection by cat breeders and cat show judges will ultimately control the health and quality of life of domestic cats with dwarfism. More detailed studies demonstrating the morphological variation should further enlighten the concerns for impaired ambulation and secondary osteoarthritis in the dwarf cat breed. Regulatory and structural variant analyses are required to further elucidate the variant causing disproportionate dwarfism in the cat.

## Acknowledgements

We appreciate the enthusiasm and provision of samples by cat breeders, especially Ms. Terri Harris. We appreciate the technical assistance of Jennifer Grahn, and Nicholas Gustafson.

## Author contributions

- Conception and design – LAL, DBF, SP
- Provision of study materials or patients – LAL, RAG, BG, MJH, NAV
- Collection and assembly of data – LAL, RAG, MJH, BG, RMB, KLC, SS, NAV
- Analysis and interpretation of the data – KLC, RAG, MJH, BG, RMB, JRC, DBF
- Drafting of the article – LAL, RMB
- Statistical expertise – JRM, RMB
- Obtaining of funding – LAL, KLC
- Critical revision of the article for important intellectual content – LAL, RMB
- Final approval of the article – all authors except SP.

## Studies involving humans or animals

The procedures followed were in accordance with the ethical standards of the responsible committee on human experimentation (institutional and national) and with the Helsinki Declaration of 1975, as revised in 2000. All sample collection and cat studies were conducted in accordance with an approved University of California, Davis Institutional Animal Care and Use protocols 11977, 15117, and 16691 and University of Missouri protocols 7808 and 8292.

---

**Supplementary Table 1.** Variants identified by whole genome sequencing within the critical region identified by genome-wide association study.

| Chromosome:<br>Position | Reference/<br>Alternative | Gene | Gene Position | Transcript Name | HGVS c. (Clinically Relevant) |
| --- | --- | --- | --- | --- | --- |
| **B1:167386513** |  | **FCA149** | **intergenic_STR** |  |  |
| B1:170278183 | C/T |  | intergenic_variant |  |  |
| **B1:170786914** |  |  | intergenic_variant |  |  |
| B1:171215074 | T/A |  | intergenic_variant |  |  |
| B1:171486710 | G/A |  | intergenic_variant |  |  |
| B1:171524991 | C/T |  | intergenic_variant |  |  |
| B1:172038327 | A/C |  | intergenic_variant |  |  |
| B1:172906886 | A/G |  | intergenic_variant |  |  |
| B1:173097092 | T/C | LIMCH1 | intron_variant | ENSFCAT00000047977 | ENSFCAT00000047977:c.237+270A>G |
| B1:173099638 | -/AAATA | LIMCH1 | intron_variant | ENSFCAT00000047977 | ENSFCAT00000047977:c.168-2211_168-2207dupTATTT |
| B1:173103057 | T/A | LIMCH1 | intron_variant | ENSFCAT00000047977 | ENSFCAT00000047977:c.168-5626A>T |
| B1:173137458 | -/T | LIMCH1 | intron_variant | ENSFCAT00000047977 | ENSFCAT00000047977:c.97-16755dupA |
| B1:173285632 | G/A |  | intergenic_variant |  |  |
| B1:173391984 | -/A | APBB2 | intron_variant | ENSFCAT00000022248 | ENSFCAT00000022248:c.-177+27299dupA |
| B1:173436205 | T/G | APBB2 | intron_variant | ENSFCAT00000022248 | ENSFCAT00000022248:c.-176-44053T>G |
| B1:173497493 | A/C | APBB2 | intron_variant | ENSFCAT00000022248 | ENSFCAT00000022248:c.-70-13022A>C |
| B1:173615935 | AGAA/- | APBB2 | intron_variant | ENSFCAT00000022248 | ENSFCAT00000022248:c.863-8872_863-8869delAAAG |
| B1:173848028 | T/G |  | intergenic_variant |  |  |
| B1:173936171 | G/T | RBM47 | intron_variant | ENSFCAT00000012401 | ENSFCAT00000012401:c.-32+31359G>T |
| B1:173949358 | TTTAT/- | RBM47 | intron_variant | ENSFCAT00000012401 | ENSFCAT00000012401:c.-32+44578_-<br>32+44582delTATTT |
| B1:173949362 | TTTTA/- | RBM47 | intron_variant | ENSFCAT00000012401 | ENSFCAT00000012401:c.-32+44578_-<br>32+44582delTATTT |
| B1:173951124 | G/A | RBM47 | intron_variant | ENSFCAT00000012401 | ENSFCAT00000012401:c.-32+46312G>A |
| B1:173970209 | C/T | RBM47 | intron_variant | ENSFCAT00000012401 | ENSFCAT00000012401:c.-32+65397C>T |
| B1:173978144 | G/C | RBM47 | intron_variant | ENSFCAT00000012401 | ENSFCAT00000012401:c.-32+73332G>C |
| B1:173978205 | C/T | RBM47 | intron_variant | ENSFCAT00000012401 | ENSFCAT00000012401:c.-32+73393C>T |
| B1:173980454 | A/G | RBM47 | intron_variant | ENSFCAT00000012401 | ENSFCAT00000012401:c.-31-74347A>G |
| B1:173990664 | T/G | RBM47 | intron_variant | ENSFCAT00000012401 | ENSFCAT00000012401:c.-31-64137T>G |
| B1:174053810 | C/A | RBM47 | intron_variant | ENSFCAT00000012401 | ENSFCAT00000012401:c.-31-991C>A |
| B1:174071994 | C/T |  | intergenic_variant |  |  |
| B1:174076407 | C/A |  | intergenic_variant |  |  |
| B1:174077557 | G/A |  | intergenic_variant |  |  |
| B1:174077762 | G/A |  | intergenic_variant |  |  |
| B1:174078299 | A/G |  | intergenic_variant |  |  |
| B1:174079184 | GAGA/- |  | intergenic_variant |  |  |
| B1:174079207 | C/A |  | intergenic_variant |  |  |
| B1:174079731 | A/G |  | intergenic_variant |  |  |
| B1:174080146 | AGAG/- |  | intergenic_variant |  |  |
| B1:174080721 | C/T |  | intergenic_variant |  |  |
| B1:174081820 | G/C |  | intergenic_variant |  |  |
| B1:174081969 | T/A |  | intergenic_variant |  |  |
| B1:174082694 | T/G |  | intergenic_variant |  |  |
| B1:174084958 | A/G |  | intergenic_variant |  |  |
| B1:174085059 | G/A |  | intergenic_variant |  |  |
| B1:174085166 | CT/- |  | intergenic_variant |  |  |
| B1:174085855 | C/T |  | intergenic_variant |  |  |
| B1:174087078 | C/T |  | intergenic_variant |  |  |
| B1:174089147 | T/C |  | intergenic_variant |  |  |
| B1:174096760 | A/T |  | intergenic_variant |  |  |
| B1:174105763 | C/T |  | intergenic_variant |  |  |
| B1:174127194 | AGTC/- | CHRNA9 | intron_variant | ENSFCAT00000042269 | ENSFCAT00000042269:c.366-145_366-142delGACT |
| B1:174131921 | G/A | CHRNA9 | intron_variant | ENSFCAT00000042269 | ENSFCAT00000042269:c.365+2136C>T |
| B1:174132207 | A/C | CHRNA9 | intron_variant | ENSFCAT00000042269 | ENSFCAT00000042269:c.365+1850T>G |
| B1:174132368 | G/A | CHRNA9 | intron_variant | ENSFCAT00000042269 | ENSFCAT00000042269:c.365+1689C>T |
| B1:174133775 | G/C | CHRNA9 | intron_variant | ENSFCAT00000042269 | ENSFCAT00000042269:c.365+282C>G |
| **B1:174566680** |  | **FCA827** | **intergenic_STR** |  |  |
| B1:174847189 | G/T |  | intergenic_variant |  |  |
| B1:174847190 | A/C |  | intergenic_variant |  |  |
| B1:174860626 | T/G | UGDH | intron_variant | ENSFCAT00000009602 | ENSFCAT00000009602:c.-7-1593T>G |
| B1:174867022 | T/- | UGDH | intron_variant | ENSFCAT00000009602 | ENSFCAT00000009602:c.163-439delT |
| B1:174999000 | A/T | RFC1 | intron_variant | ENSFCAT00000001781 | ENSFCAT00000001781:c.135+5393A>T |
| B1:175078887 | A/G | WDR19 | intron_variant | ENSFCAT00000001779 | ENSFCAT00000001779:c.*13+3682T>C |
| B1:175078920 | A/G | WDR19 | intron_variant | ENSFCAT00000001779 | ENSFCAT00000001779:c.*13+3649T>C |
| B1:175078933 | T/C | WDR19 | intron_variant | ENSFCAT00000001779 | ENSFCAT00000001779:c.*13+3636A>G |
| B1:175078947 | C/T | WDR19 | intron_variant | ENSFCAT00000001779 | ENSFCAT00000001779:c.*13+3622G>A |
| B1:175078953 | T/C | WDR19 | intron_variant | ENSFCAT00000001779 | ENSFCAT00000001779:c.*13+3616A>G |
| B1:175349169 | A/G | TMEM156 | intron_variant | ENSFCAT00000031918 | ENSFCAT00000031918:c.89-5273A>G |
| B1:175975857 | G/C |  | intergenic_variant |  |  |
| **B1:179961345** |  | **FCA152** | **intergenic_STR** |  |  |
| **B1:186605881** |  |  | intergenic_variant |  |  |

**Supplementary Table 2.** Critical region genes for feline dwarfism on cat chromosome B1:170278183-175975857.

| Gene ID | Name | Start | End | GeneCard associated Diseases |
| --- | --- | --- | --- | --- |
| ENSFCAG00000028486 | *GNPDA2* | 170293762 | 170322496 | Obesity |
| ENSFCAG00000028580 | *GUF1* | 170325353 | 170346889 | Epileptic Encephalopathy, West Syndrome |
| ENSFCAG00000006282 | *YIPF7* | 170366863 | 170394543 | De Quervain Disease, Epicondylitis. |
| ENSFCAG00000029885 |  | 170488906 | 170489280 |  |
| ENSFCAG00000030570 | *KCTD8* | 170589687 | 170845851 | Sweet Taste Signaling, Neuropathic Pain-Signaling in Dorsal Horn Neurons |
| ENSFCAG00000034150 | *RF00100* | 171550935 | 171551217 |  |
| ENSFCAG00000018569 | *GRXCR1* | 171751919 | 171878987 | Deafness, Branchiootic Syndrome 1 |
| ENSFCAG00000036356 |  | 171990790 | 171991881 |  |
| ENSFCAG00000015213 | *ATP8A1* | 172062098 | 172289239 | Cerebellar Ataxia, Mental Retardation, Dysequilibrium Syndrome, Robinow Syndrome |
| ENSFCAG00000043473 | *SHISA3* | 172299898 | 172305862 | Tumor suppressor |
| ENSFCAG00000033789 | *RF00026* | 172345184 | 172345284 |  |
| ENSFCAG00000044789 |  | 172396346 | 172417546 |  |
| ENSFCAG00000043262 |  | 172442984 | 172449907 |  |
| ENSFCAG00000044080 | *BEND4* | 172498442 | 172530596 | Follicular Lymphoma |
| ENSFCAG00000024533 | *SLC30A9* | 172564833 | 172650293 | Birk-Landau-Perez Syndrome |
| ENSFCAG00000040948 | *TMEM33* | 172673293 | 172693525 | Congenital Ichthyosis |
| ENSFCAG00000045230 | *PHOX2B* | 172854075 | 172859113 | Central Hypoventilation Syndrome, Congenital Neuroblastoma 2. |
| ENSFCAG00000042497 |  | 172866353 | 172866487 |  |
| ENSFCAG00000046477 |  | 172869997 | 172882934 |  |
| ENSFCAG00000007444 | *LIMCH1* | 172914742 | 173249269 | Lung cancer, Hepatosplenic T-Cell Lymphoma |
| ENSFCAG00000025457 | *UCHL1* | 173311731 | 173327417 | Parkinson disease |
| ENSFCAG00000012843 | *APBB2* | 173364437 | 173753735 | Alzheimer Disease |
| ENSFCAG00000012841 | *NSUN7* | 173754945 | 173809607 | Infertility |
| ENSFCAG00000042869 |  | 173865395 | 173885964 |  |
| ENSFCAG00000012399 | *RBM47* | 173904131 | 174069382 | Breast, Colorectal Cancer |
| ENSFCAG00000035547 | *CHRNA9* | 174118937 | 174135747 | Deafness, Breast cancer |
| ENSFCAG00000012394 | *RHOH* | 174208322 | 174208897 | T-Cell Immunodeficiency with Epidermodysplasia Verruciformis |
| ENSFCAG00000024887 | *N4BP2* | 174283177 | 174356646 |  |
| ENSFCAG00000037355 | *RF00026* | 174304986 | 174305092 |  |
| ENSFCAG00000003209 | *PDS5A* | 174441923 | 174579837 |  |
| FCA827 | *FCA827* | 174566680 | 174566701 |  |
| ENSFCAG00000033193 | *UBE2K* | 174615267 | 174689895 | Huntington Disease |
| ENSFCAG00000038568 | *RF00007* | 174675118 | 174675271 |  |
| ENSFCAG00000024033 |  | 174747387 | 174747617 |  |
| ENSFCAG00000032694 | *SMIM14* | 174760874 | 174841113 |  |
| ENSFCAG00000009600 | *UGDH* | 174853638 | 174885557 | Deafness, biosynthesis of glycosaminoglycans such as hyaluronan, chondroitin sulfate, and heparan sulfate |
| ENSFCAG00000023567 | *LIAS* | 174892614 | 174907965 | Hyperglycinemia, Lactic Acidosis, Seizures, Lipoic Acid Synthetase Deficiency. |
| ENSFCAG00000012284 | *RPL9* | 174908042 | 174912983 | Lipoic Acid Synthetase Deficiency. |
| ENSFCAG00000026771 | *KLB* | 174918899 | 174953922 | Hypogonadotropic Hypogonadism |
| **ENSFCAG00000001781** | ***RFC1*** | **174993455** | **175063047** | **Hutchinson-Gilford Progeria Syndrome** |
| ENSFCAG00000001779 | *WDR19* | 175062084 | 175177435 | Disorders affecting function of the cilium |
| ENSFCAG00000003534 | *KLHL5* | 175208337 | 175296713 |  |
| ENSFCAG00000028887 | *TMEM156* | 175327852 | 175381406 | Follicular Lymphoma |
| ENSFCAG00000027380 | *FAM114A1* | 175397428 | 175467463 | Mixed Germ Cell Cancer |
| ENSFCAG00000043678 | *RF00026* | 175459211 | 175459317 |  |
| ENSFCAG00000040731 |  | 175493242 | 175495632 |  |
| ENSFCAG00000031748 |  | 175521624 | 175523997 |  |
| ENSFCAG00000007666 | *TLR10* | 175549356 | 175551785 | Crimean-Congo Hemorrhagic Fever, Theileriasis. |
| ENSFCAG00000028338 | *KLF3* | 175613723 | 175648997 | Transcriptional misregulation in cancer |

